# Direct readout of heterochromatic H3K9me3 regulates DNMT1-mediated maintenance DNA methylation

**DOI:** 10.1101/2020.04.27.064493

**Authors:** Wendan Ren, Huitao Fan, Sara A Grimm, Yiran Guo, Jae Jin Kim, Linhui Li, Christopher James Petell, Xiao-Feng Tan, Zhi-Min Zhang, John P. Coan, Jiekai Yin, Linfeng Gao, Ling Cai, Brittany Detrick, Burak Çetin, Yinsheng Wang, Qiang Cui, Brian D. Strahl, Or Gozani, Kyle M. Miller, Seán E. O’Leary, Paul A. Wade, Dinshaw J. Patel, Gang Greg Wang, Jikui Song

**Author notes:** School of pharmacy, Jinan University, 601 Huangpu Avenue West, Guangzhou 510632, China (Zhi-Min Zhang). These authors contributed equally to this work (Wendan Ren, Huitao Fan).

## Abstract

In mammals, repressive histone modifications such as trimethylation of histone H3 Lys9 (H3K9me3), frequently coexist with DNA methylation, producing a more stable and silenced chromatin state. However, it remains elusive how these epigenetic modifications crosstalk. Here, through structural and biochemical characterizations, we identified the replication foci targeting sequence (RFTS) domain of maintenance DNA methyltransferase DNMT1, a module known to bind the ubiquitylated H3 (H3Ub), as a specific reader for H3K9me3/H3Ub, with the recognition mode distinct from the typical trimethyl-lysine reader. Disruption of the interaction between RFTS and the H3K9me3Ub affects the localization of DNMT1 in stem cells and profoundly impairs the global DNA methylation and genomic stability. Together, this study reveals a previously unappreciated pathway through which H3K9me3 directly reinforces DNMT1-mediated maintenance DNA methylation.

## Introduction

DNA methylation is an evolutionarily conserved epigenetic mechanism that critically influences chromatin structure and function (1). In mammals, DNA methylation predominantly occurs at the C-5 position of cytosine within the CpG dinucleotide context, which regulates the silencing of retrotransposons (2), allele-specific genomic imprinting (3), X-chromosome inactivation (4), and tissue-specific gene expression that underlies cell fate commitment (5). During DNA replication, DNA methylation is stably propagated by DNA methyltransferase 1 (DNMT1), which replicates the DNA methylation patterns from parental DNA strands to the newly synthesized strands in a replication-dependent manner (6, 7). Faithful propagation of DNA methylation patterns is essential for clonal transmission of epigenetic regulation between cell generations.

DNMT1 is a multi-domain protein, comprised of a large N-terminal regulatory region and a C-terminal methyltransferase (MTase) domain, linked via a conserved (GK)n dipeptide repeat (Fig. 1A). The regulatory region contains a replication-foci-targeting sequence (RFTS), a CXXC zinc finger domain and a pair of bromo-adjacent homology (BAH) domains (8–11). Recent structural and biochemical evidence has revealed that both the RFTS and CXXC domains regulate the activity of DNMT1 through autoinhibitory mechanisms: the RFTS domain directly interacts with the MTase domain to inhibit DNA binding (10–14) whereas the CXXC domain specifically recognizes unmethylated CpG nucleotides, which in turn blocks the *de novo* methylation activity of DNMT1 (8, 15). These N-terminal domains-mediated allosteric regulations, together with an inherent enzymatic preference of the MTase domain for hemimethylated CpG sites (9, 16), shape the enzymatic specificity of DNMT1 in maintaining DNA methylation.

**Figure 1.**
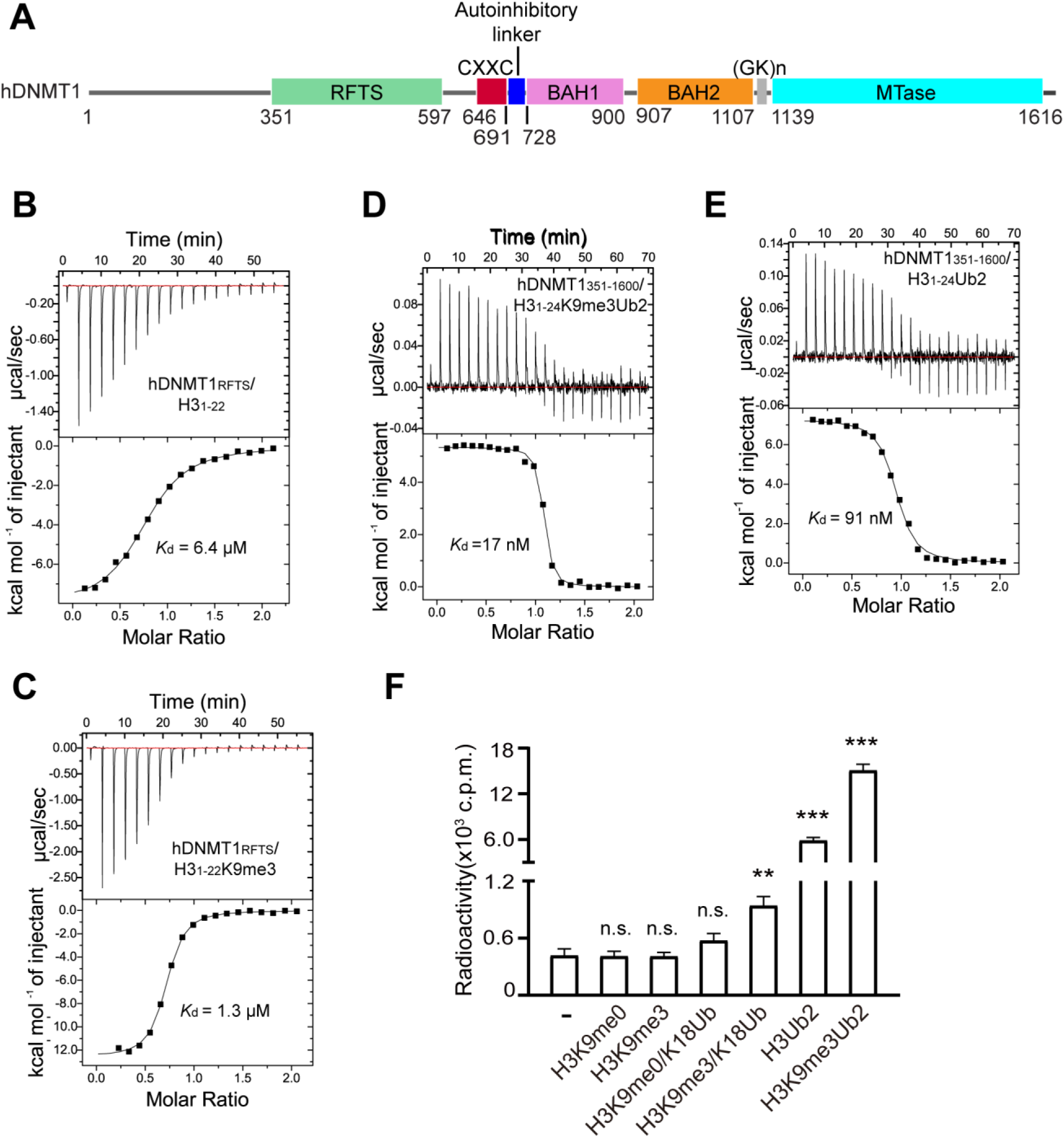
Specific interaction between the DNMT1 RFTS domain and H3K9me3Ub. (**A**) Domain architecture of human DNMT1 (hDNMT1), with individual domains delimited by residues numbers. (**B,C**) ITC binding assays of hDNMT1_RFTS_ over H3_1-22_ (**B**) and H3_1-22_K9m3 (**C**) peptides. (**D,E**) ITC binding assays of hDNMT1_351-1600_ over H3_1-24_K9me3Ub2 (**D**) and H3_1-24_Ub2 (**E**) peptides. (**F**) *In vitro* DNA methylation assays for hDNMT1_351-1600_ in the absence or presence of H3 peptides with the indicated modification. Mean and s.d. were derived from three independent measurements. n.s., not significant; \*\**p* < 0.01; \*\*\**p* < 0.001.

The crosstalk between DNA methylation and other gene silencing mechanisms, such as histone H3K9 methylation, is widely observed across evolution, ensuring proper chromatin assembly and loci-specific gene suppression (17, 18). The H3K9 methylation-dependent DNA methylation in *Neurospora* and *Arabidopsis* has been elucidated, which is attributed to a direct or indirect H3K9me2/H3K9me3 readout mechanism of DNA methyltransferases (19, 20). In mammals, H3K9me3 is strongly correlated with DNA methylation at pericentric heterochromatin (21). However, the mechanism by which H3K9 methylation is translated into mammalian DNA methylation remains far from being fully understood. Nevertheless, it has been established that DNMT1-mediated maintenance DNA methylation is critically regulated by Ubiquitin-like, containing PHD and RING Finger domains, 1 (UHRF1) (22, 23). During the S phase, UHRF1 is recruited to replicating heterochromatin through its association with both hemi-methylated CpG DNA and H3K9me3 (24), where it stochastically catalyzes the mono-ubiquitylation of histone H3 at lysine 14 (H3K14Ub), lysine 18 (H3K18Ub) and/or lysine 23 (H3K23Ub) (25–28), and PCNA-associated factor 15 (PAF15) at lysine 15 and 24 (29, 30). The DNMT1 RFTS domain recognizes all these modifications, with a preference for the two-mono-ubiquitin marks (i.e. H3K18Ub/H3K23Ub), leading to allosteric stimulation of DNMT1 (28). The structure of DNMT1 RFTS domain in complex with H3K18Ub/K23Ub revealed that the H3K18Ub/K23Ub binding leads to structural rearrangement of RFTS and its dissociation with the C-terminal linker, thereby facilitating the conformation transition of DNMT1 from an autoinhibitory state to an active one (28, 31). These observations suggest a UHRF1-bridged link between H3K9me3 and DNA methylation. However, whether H3K9me3 directly interacts with DNMT1 to regulate maintenance DNA methylation remains unknown.

To determine how H3K9me3 affects DNMT1-mediated maintenance DNA methylation, we examined the histone binding activity of DNMT1 RFTS domain and its relationship to the chromatin association and enzymatic activity of DNMT1. Through Isothermal Titration Calorimetry (ITC) and *in vitro* enzymatic assays, we identified that the RFTS domain binds preferably to H3K9me3 over H3K9me0, which serves to strengthen the enzymatic stimulation of DNMT1 by H3 ubiquitylation (H3Ub). Furthermore, we determined the crystal structure of bovine RFTS domain complexed with H3K9me3 peptide and two ubiquitins, providing the molecular basis for the H3K9me3 recognition. In addition, our cellular and genomic methylation analysis demonstrated that impairment of the RFTS-H3K9me3Ub recognition led to reduced co-localization of DNMT1 with H3K9me3, a global loss of DNA methylation patterns and genome instability in mouse embryonic stem (ES) cells. Together, this study provides a mechanism by which H3K9me3 directly regulates DNMT1-mediated maintenance DNA methylation in mammalian cells.

## Results

### The RFTS domain of DNMT1 is an H3K9me3Ub reader module

Recent studies have indicated that UHRF1-mediated recognition of H3K9me3 and hemi-methylated CpG DNA stimulates its E3 ubiquitin ligase activity on histone H3 (27, 32), thus providing a linkage between H3K9me3 and H3 ubiquitylation during UHRF1/DNMT1-mediated DNA methylation. This observation prompted us to ask whether or not H3K9me3 directly influences the interaction between the RFTS domain of DNMT1 and ubiquitylated H3. To this end, we performed ITC assays using the purified human DNMT1 RFTS domain (hDNMT1_RFTS_) and histone H3 peptides (residues 1-22, H3_1-22_), either unmodified or with H3K9me3 modification (H3_1-22_K9me3) (Table S1). Titration of hDNMT1_RFTS_ with H3_1-22_ gives a dissociation constant (*K_d_*) of 6.4 μM (Fig. 1B). In comparison, titration of hDNMT1_RFTS_ with H3_1-22_K9me3 gives a *K*_d_ of 1.3 μM, suggesting that hDNMT1_RFTS_ has a ~5-fold binding preference for H3K9me3 over unmodified H3 tails (Fig. 1C). Furthermore, we probed the interaction of hDNMT1_RFTS_ with the H3_1-22_ peptide acetylated at K9 (H3_1-22_K9Ac) and other histone tri-methylations (H3K4me3, H3K27me3, H3K36me3 and H4K20me3) and observed a much weaker binding (Fig. S1A-S1E). These observations not only confirm the previously observed interaction between hDNMT1_RFTS_ and H3 (28), but also support a preferential binding of H3K9me3 over H3K9me0 by hDNMT1_RFTS_.

Next, we asked whether H3K9me3 can enhance the interaction between H3K18Ub/K23Ub and the RFTS domain in the context of full-length DNMT1. Following a previously established approach (28, 33), we installed dual ubiquitin marks onto a K18C/K23C mutant form of H3_1-24_ (H3_1-24_Ub2) or H3_1-24_K9me3 (H3_1-24_K9me3Ub2) peptides through dichloroacetone (DCA) linkage (33), followed by ITC binding assay against a hDNMT1 fragment (residues 351-1600, hDNMT_351-1600_) spanning from the RFTS domain to the C-terminal MTase domain (Fig. 1A). We found that hDNMT_351-1600_ binds to the H3_1-24_K9me3Ub2 peptide with a *K*_d_ of 17 nM (Fig. 1D), exhibiting a ~5-fold binding preference over the H3_1-24_Ub2 peptide (*K_d_* = 91 nM) (Fig. 1E). These data support that, together with H3Ub, H3K9me3 provides a signal for tethering DNMT1 onto chromatin via its RFTS domain.

### The H3K9me3 readout boosts the stimulation effect of H3Ub on DNMT1 activity

In light of a previous study showing that recognition of H3Ub by the DNMT1 RFTS domain results in enhanced enzymatic activity of DNMT1 (28), we have further evaluated the effect of H3K9me3 on the enzymatic stimulation of hDNMT1. Here, we found that, consistent with the previous observation (28), incubation of hDNMT1_351-1600_ with equimolar H3K18Ub and H3Ub2 led to enhanced hDNMT1_351-1600_ activity by 1.3 and 14 fold, respectively (Fig. 1F). In contrast, incubation with equimolar H3K9me3/K18Ub and H3K9me3Ub2 promoted the enzymatic activity of hDNMT1_351-1600_ by 2.2 and 36 fold, respectively. Note that incubation of hDNMT1_351-1600_ with equimolar H3K9me0 or H3K9me3 did not change its activity appreciably (Fig. 1F), although at increased peptide concentrations, the activity of hDNMT1_351-1600_ is significantly higher when incubated with H3K9me3, relative to H3K9me0 peptide controls (Fig. S1F). Together, these results show the specific recognition of H3K9me3 over H3K9me0 by hDNMT1_RFTS_ elevates the stimulation effect of H3Ub, a transient ligand of hDNMT1_RFTS_ as previously shown (25, 28).

### The structure of RFTS domain in complex with H3K9me3 peptide and two-mono ubiquitin

To gain a molecular understanding of the RFTS-H3K9me3 recognition, we next crystallized the complex of the RFTS domain from bovine DNMT1 (bDNMT1_RFTS_) with both the H3_1-24_K9me3/K18C/K23C peptide and G76C-mutated ubiquitin, and solved the structure at 3.0 Å resolution (Fig. 2A and Table S2). The structure reveals a bDNMT1_RFTS_-H3K9me3-Ub2 complex, with clearly traced H3 peptide from Arg2 to Leu20 (Fig. 2B). The bDNMT1_RFTS_ domain folds into a two-lobe architecture, with an N-lobe harboring a zinc finger and a C-lobe dominated by a helical bundle (Fig. 2A), resembling that of hDNMT1_RFTS_ (11, 14, 28, 31). Like the previously reported hDNMT1_RFTS_-H3K18Ub/K23Ub complex (28), the H3K9me3 peptide traverses across the surface of the C-lobe and N-lobe of the bDNMT1_RFTS_ domain and engages extensive intermolecular interactions, with the side chain of H3K9me3 residue embraced by a surface groove at the interface between bDNMT1_RFTS_ and one of the ubiquitin molecules (Fig. 2B and C). The two ubiquitin molecules are mostly positioned at the N-lobe of the RFTS domain, separated by the ubiquitin recognition loop (URL; residues 386-398), and interact with the RFTS domain and H3 in a fashion similar to what was previously observed for the hDNMT1_RFTS_-H3K18Ub/K23Ub complex (Fig. 2C-F) (28). Of particular note, residue G76C of one ubiquitin and H3 K18C are positioned in a distance that allows disulfide bond formation, while the C-terminus of the other ubiquitin is near to the side chain of H3K14 and also likely accessible to the disordered H3K23 (Fig. 2G), reminiscent of the close proximity of these moieties in the hDNMT1RFTS-H3K18Ub/K23Ub complex. Note that in the hDNMT1RFTS-H3K18Ub/K23Ub complex, the H3 peptide was conjugated with two G76C-mutated ubiquitins through disulfide linkages before mixing with hDNMT1_RFTS_, whereas in the current complex, no covalent linkage was involved between the H3K9me3 peptide and the ubiquitins prior to the complex assembly. In this regard, the similar positioning of H3 and ubiquitins between the two complexes, crystallized under different conditions, reinforces the notion that H3 and ubiquitin molecules are able to engage DNMT1_RFTS_ via independent inter-molecular interactions (28). These results also indicate that H3K9me3 and H3Ub2 may act synergistically or independently for associating DNMT1 onto chromatin.

**Figure 2.**
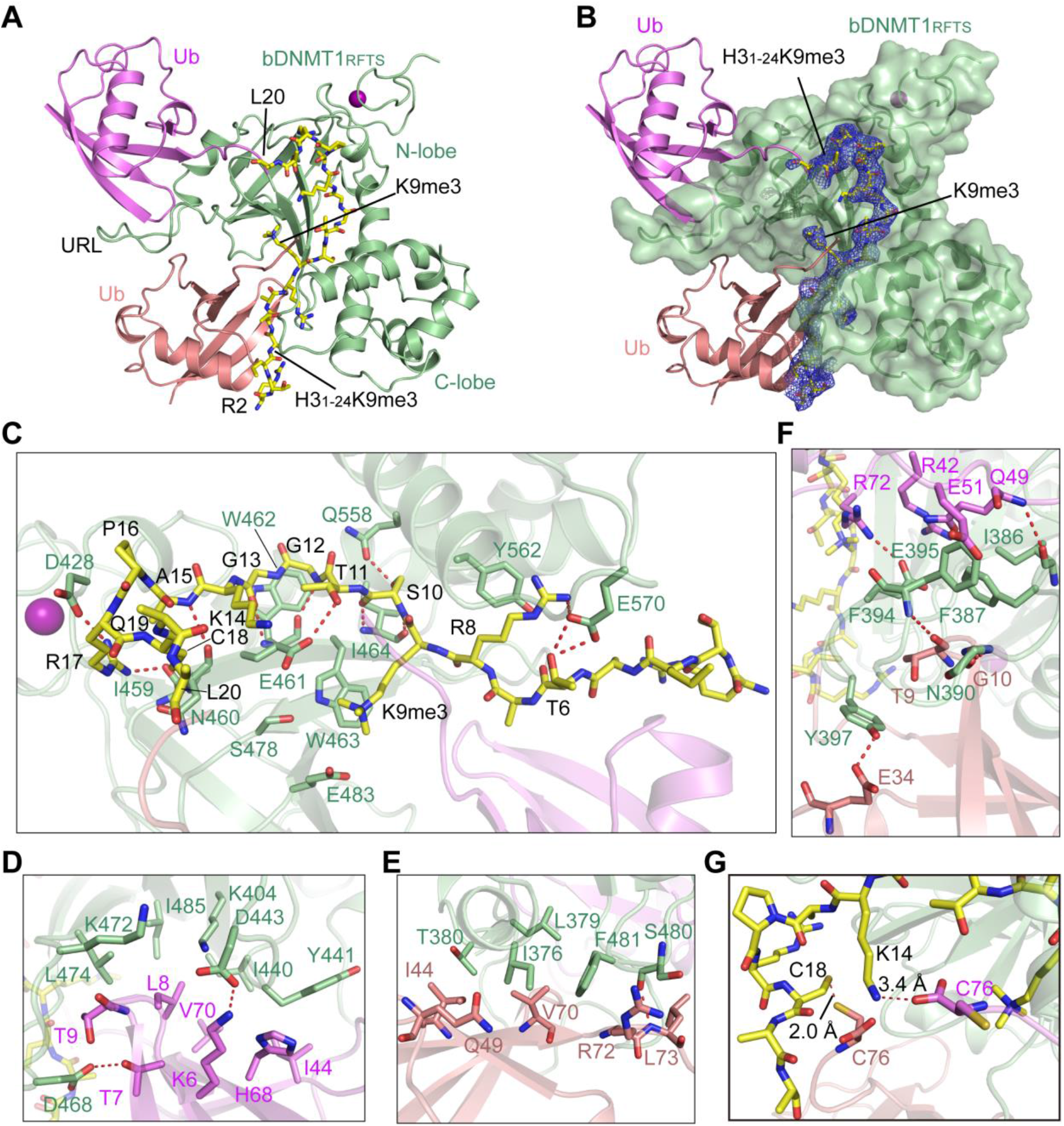
Structural details for the bDNMT1_RFTS_ domain in complex with H3K9me3K18C/K23C and ubiquitin. **(A)** Crystal structure of bovine DNMT1_RFTS_ (light green) in complex with H3_1-24_K9me3/K18C/K23C peptide (yellow) and G76C-mutated ubiquitin (magenta and salmon). The zinc ions are shown as purple spheres. (**B**) Crystal structure of bovine DNMT1_RFTS_ (light green) in complex with H3_1-24_K9me3/K18C/K23C peptide (yellow) and G76C-mutated ubiquitin (purple and salmon). The Fo-Fc omit map (blue) of the H3 peptide was contoured at the 1.5 σ level. (**C**) Close-up views of the intermolecular interactions between bDNMT1 RFTS (light green) and the H3_1-24_K9me3K18CK23C peptide (yellow stick). The two ubiquitin molecules are colored in salmon and magenta, respectively. Hydrogen bonds are shown as dashed lines. The zinc ion is shown as purple sphere. (**D,E**) Close-up views of the intermolecular interactions between the conserved I44 patch of two ubiquitin molecules (magenta and salmon) and bDNMT1 RFTS (light green). (**F**) Close-up views of the intermolecular interactions between URL of bDNMT1 RFTS (light green) and two ubiquitin molecules (magenta and salmon). (**G**)Close-up views of the close proximity between the H3_1-24_K9me3K18CK23C peptide (yellow stick) and the ubiquitin molecules (salmon and magenta).

Strikingly, in our structure, we found that both DNMT1 and H3Ub contribute to H3K9me3 engagement. First, H3K9me3 stacks its side chain against the indole ring of bDNMT1_RFTS_ W463, with the quaternary ammonium group interacting with the side chain carboxylates of bDNMT1_RFTS_ E461 and E483 through electrostatic attractions (Fig. 3A). In addition, residues R72-G75 of one of the bound ubiquitins make both main-chain and side-chain van der Waals contacts with H3K9me3 from an opposite direction, leading to formation of an H3K9me3-binding pocket (Fig. 3A). Such an H3K9me3-binding site of DNMT1_RFTS_ is distinct from the multi-walled aromatic cage typical for a trimethyl-lysine (Kme3) reader (34), but is reminiscent of the interaction between the PHD finger domain of E3 ubiquitin ligase TRIM33 and H3K9me3 (Fig. S2A) (35). Through superimposition with the previously determined hDNTM1_RFTS_-H3K18Ub/K23Ub structure (28), we found that those two structures overlap very well, with a root-mean-square deviation (RMSD) of 0.6 Å over 384 aligned Cα atoms (Fig. S2B). However, in comparison to the H3K9me0 binding, association of H3K9me3 with bDNMT1_RFTS_ W463 leads to an increase in buried surface area of the H3K9-engaging pocket by 8-9 Å^2^ (Fig. S2B).

**Figure 3.**
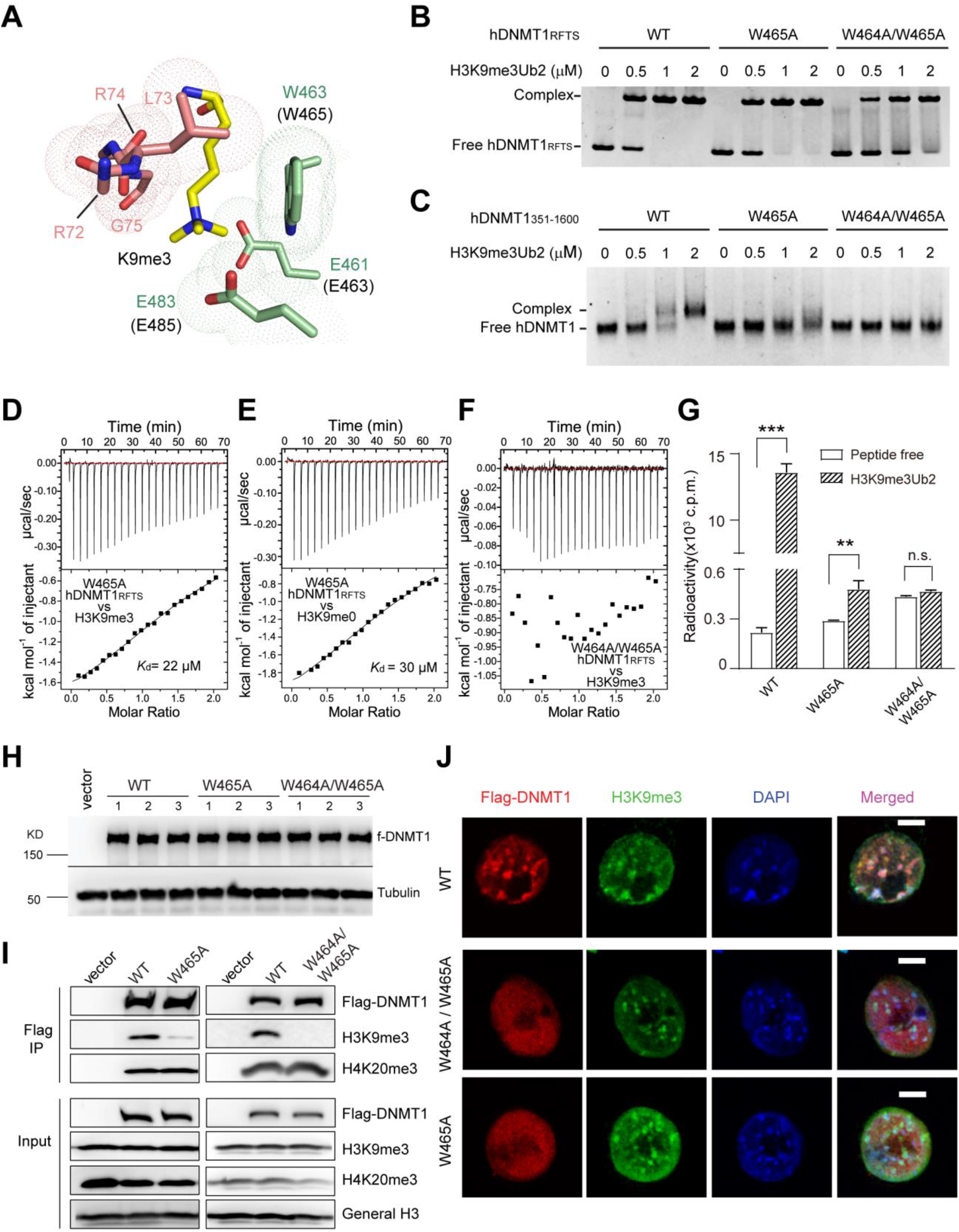
Biochemical and cellular analysis of the RFTS-H3K9me3Ub2 interaction. (**A**) Close-up view of the residues forming the H3K9me3-binding pocket, in the same color scheme as in Fig. 2. For clarity, the side chains of ubiquitin R72 and R74 are not shown. (**B**) EMSA analysis of the interaction between hDNMT1_351-597_, either wild type (WT), W465A or W464A/W465A, and the H3_1-24_K9me3Ub2 peptide. (**C**) EMSA analysis of the interaction between hDNMT1_351-1600_, either wild type (WT), W465A or W464A/W465A, and the H3_1-24_K9me3Ub2 peptide. (**D,E**) ITC binding assays of hDNMT1_RFTS_ W465A mutant over H3_1-22_K9m3 (**D**) and H3_1-22_ (**E**) peptides. (**F**) ITC binding assays of hDNMT1_RFTS_ W464A/W465A mutant over H3_1-22_K9me3 peptide. (**G**) Immunoblots of the indicated Flag-tagged DNMT1 after stable reconstitution into the independently derived 1KO-ESC lines. (**H**) CoIP (upper panels) detecting association of the indicated Flag-tagged DNMT1 with H3K9me3 or H4K20me3. Bottom panels are immunoblots of input. (**I**) DNA methylation activities of hDNMT1_351-1600_, either WT, W465A or W464A/W465A, in the absence of presence of H3_1-24_K9me3Ub2 peptide. Mean and s.d. were derived from three independent measurements. (**, *p* < 0.01, ***, *p* < 0.001, n.s. not significant, Student’s *t*-test) (**J**) Representative confocal immunofluorescence images revealing localization of the indicated DNMT1 (Flag-tagged, red), H3K9me3 (green) and chromatin (stained by DAPI, blue) in the 1KO-ESC stable expression lines synchronized at S phase. Scale bar, 5 micrometers.

Sequence analysis of the DNMT1 RFTS domains reveals that both the H3K9me3- and ubiquitin-binding sites are highly conserved across evolution (Fig. S2C). Among these, a di-tryptophan motif (W462 and W463 in bDNMT1) is unvaried from zebrafish to human (Fig. S2C), in line with their important roles in binding to H3K9me3 and surrounding H3 tail residues (Fig. 2C and 3A).

### Mutational analysis of the DNMT1-H3K9me3Ub2 binding

To test the above structural observations, we selected hDNMT1 W464 and W465, corresponding to bDNMT1 W462 and W463, respectively (Fig. S2C), for mutagenesis. Using Electrophoretic Mobility Shift Assays (EMSA), we found that the mutation of hDNMT1 W465 into alanine (W465A) leads to a significant reduction in the H3K9me3Ub2-binding affinity of hDNMT1_RFTS_ and hDNMT1_351-1600_ (Fig. 3B and C), while the W464A/W465A mutation reduced the hDNMT1RFTS-H3K9me3Ub2 binding even further (Fig. 3B) and led to nearly undetectable binding between hDNMT1_351-1600_ and H3K9me3Ub2 (Fig. 3C). Likewise, the ITC assays indicated that the W465A mutation reduced the H3K9me0- and H3K9me3-binding affinities of hDNMT1RFTS by ~5 and ~17 fold, respectively (Fig. 3D and E); introduction of a W464A/W465A mutation largely abolished the hDNMT1RFTS-H3K9me3 binding (Fig. 3F). Furthermore, we performed *in vitro* DNA methylation assays to evaluate the effect of these mutations on the enzymatic activity of DNMT1. Unlike wild-type hDNMT1351-1600, which shows substantial increase in methylation efficiency in the presence of H3K9me3K18Ub and H3K9me3Ub2 (Fig. 1F) as well as an enzymatic preference for H3K9me3 over H3K9me0 (Fig. S1F), introduction of the W465A or W464A/W465A mutation into hDNMT1351-1600 greatly dampened the methylation-stimulating effects of H3K9me3, H3K9me3K18Ub or H3K9me3Ub2 (Fig. 3G and Fig. S3A and B). These data support the important roles of the W464 and W465 residues in mediating the DNMT1-H3K9me3/H3Ub interaction and the consequent enzymatic activation of DNMT1.

Given that H3Ub is reportedly a transient mark in cells (36), we have focused on examining the requirement of the DNMT1_RFTS_ domain for H3K9me3 binding in cells. To this end, we used the Dnmt1-knockout mouse embryonic stem cells (1KO-ESC) (37) and generated multiple 1KO-ESC lines with comparable, stable expression of exogenous DNMT1, either wild-type (DNMT1^WT^) or an H3K9me3-binding-defective mutant (DNMT1^W465A^ or DNMT1^W464A/W465A^) (Fig. 3H). By co-immunoprecipitation (CoIP), we found that both the W464A and W464A/W465A mutations interfere with efficient binding of DNMT1 to H3K9me3, but not H4K20me3, in cells (Fig. 3I). Furthermore, in the ES cells synchronized at S phase, confocal immunofluorescence (IF) microscopy showed punctate nuclear foci of DNMT1^WT^ that overlap with the H3K9me3-marked, DAPI-dense regions of chromatin (Fig. 3J), whereas the RFTS-mutant DNMT1 proteins show a more diffuse distribution in the nucleus and lose their co-localization with H3K9me3 (Fig. 3J). Together, these data support that the histone-engaging activity of the RFTS domain is critical for both the enzymatic stimulation and chromatin occupancy of DNMT1.

### The RFTS domain of DNMT1 is crucial for global DNA methylation in cells

Next, we sought to examine the role of the DNMT1 RFTS domain in maintenance DNA methylation in cells. First, we used liquid chromatography-mass spectrometry (LC-MS) to quantify global levels of methylated cytosine (5mC) in 1KO-ESC cells reconstituted with DNMT1^WT^ or the RFTS-defective mutant. As expected, stable transduction of DNMT1^WT^ led to an increase in overall 5mC level (Fig. 4A, WT vs. 1KO). However, such an increase was found compromised by the W465A or W464A/W465A mutation, with the latter exhibiting a more severe DNA methylation defect, lacking significant methylation-stimulating effect relative to mock (Fig. 4A). Further, we carried out genome-wide methylation profiling with enhanced reduced representation bisulfite sequencing (eRRBS). Our eRRBS data showed the desired bisulfite conversion rates (Fig. S4A, 99.79-99.83%; and Table S3), with at least 5-fold coverage for ~5-8 million of CpG sites in all samples. Relative to 1KO-ESCs reconstituted with DNMT1^WT^, those with DNMT1^W465A^ or DNMT1^W464A/W465A^ showed a marked decrease in overall CpG methylation, with a complete loss of most heavily methylated CpG sites that are typically seen at heterochromatic repetitive elements (38) (Fig. 4B, red; and Fig. S4B and C). In particular, there is a significantly decreased level of CpG methylation at the H3K9me3-decorated genomic regions in cells reconstituted with either DNMT1^W465A^ or DNMT1^W464A/W465A^, relative to DNMT1^WT^ controls (Fig. 4C), as demonstrated by sub-telomeric regions located in the chromosomes 1 and X (Fig. 4D and Fig. S4D). Again, these cellular assays show that the double RFTS mutant (DNMT1^W464A/W465A^) produced more severe CpG methylation defects, relative to DNMT1^W465A^ (Fig. 4A-D and Fig. S4A-D), which is consistent with our *in vitro* biochemical and enzymatic observations. Collectively, we have demonstrated an essential role of the histone-engaging RFTS domain in maintenance of CpG methylation at the H3K9me3-associated heterochromatin in cells.

**Figure 4.**
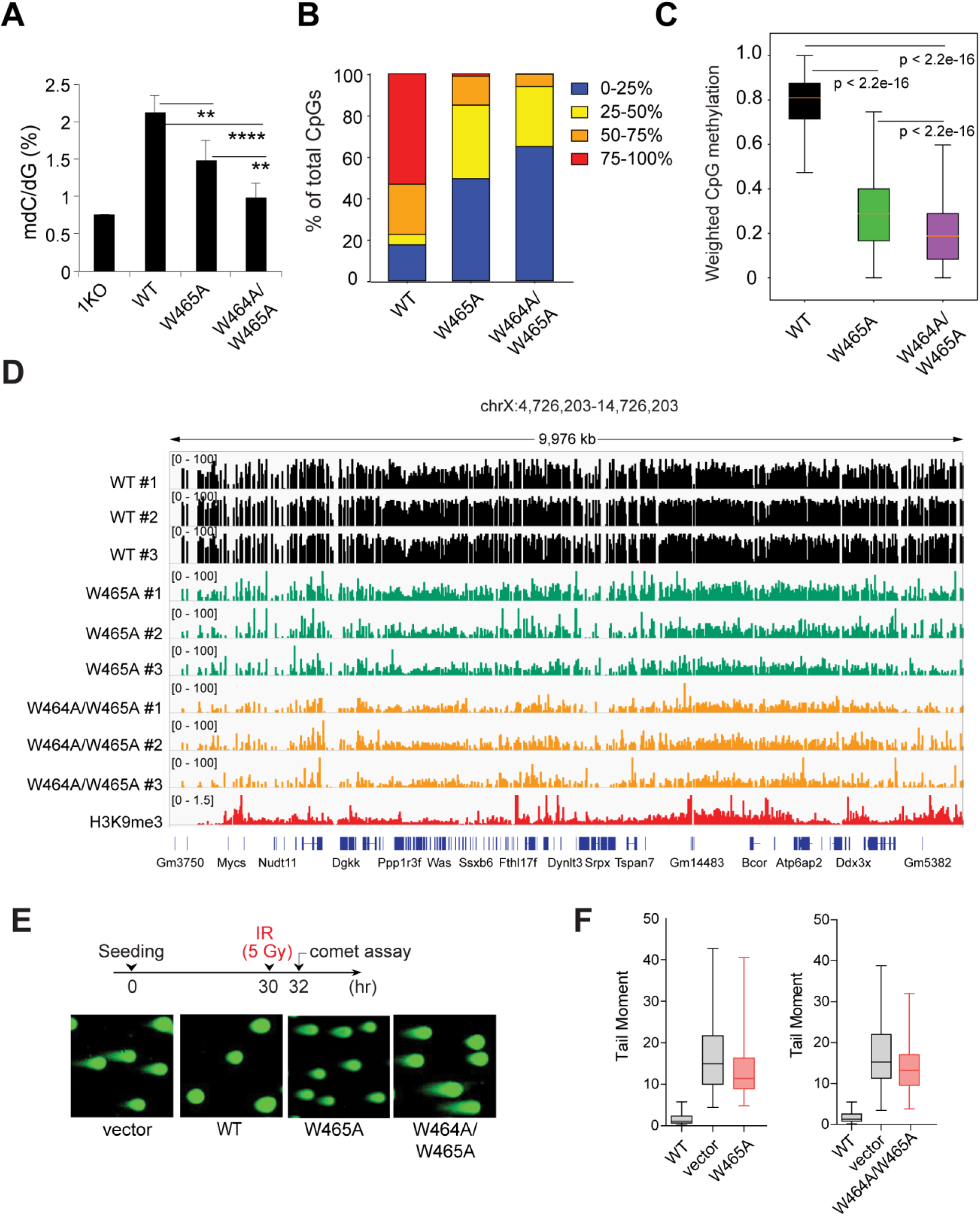
The role of RFTS mutant in cellular CpG methylation pattern and IR response. (**A**) LC-MS analysis of global 5-methyl-2-deoxycytidine (5-mdC) levels (calculated as 5-mdC/2-deoxyguanosine on the y-axis) in 1KO-ESC lines after stable transduction of empty vector or the indicated DNMT1 (n = 3-6 biological replicates). Data are mean± s.d. (**B**) Bar plots showing the CpG methylation levels in 1KO-ESC lines with stable expression of the indicated DNMT1. (**C**) Box plot shows weighted methylation levels of CpGs located within H3K9me3 peaks in 1KO-ESC lines with stable expression of the indicated DNMT1. Scores are calculated after aggregating data from three replicated samples per group. Only H3K9me3 peaks with the mapped reads are included (n = 103,603 for WT, 101,791 for W465, and 102,892 for W464A/W465A). In the plot, the box depicts the 25th to 75th percentiles, with the band in the box representing the median. (**D**) Representative IGV view shows CpG methylation at an H3K9me3-marked genomic region located in the chromosome X among three replicated 1KO-ESC lines with stable expression of the indicated DNMT1. Cytosines covered by at least 5 reads according to eRRBS data are shown, with each site designated by a vertical line. (**E,F**) Neutral comet assays revealing DNA breaks (DNA breaks quantified in panel **F**) after ionizing radiation (IR) treatment of 1KO-ESC cells reconstituted with vector control or the indicated DNMT1. Box-and whisker plots in panel F depict 25-75% in the box, whiskers are 10-90%, and median is indicated. Data represent the mean ± S.E.M. from >100 cells (N = 3 biologically independent replicates).

Maintenance of proper DNA methylation levels in cells is crucial for genome stability (39). To investigate the role of DNMT1-mediated DNA methylation in genome stabilization, we further challenged 1KO-ESC cells, reconstituted with WT or mutant DNMT1, with ionizing radiation (IR) treatment. By using the neutral comet assay (Fig. 4E), a surrogate method for scoring DNA double-strand break (DSB) lesions, we found that loss of DNMT1 rendered ESC cells a hyper-sensitivity to IR treatment, reflecting a possible change in chromatin structure or impairment in DSB repair (Fig. 4E and F), a phenotype that can be rescued by complementation with WT DNMT1. Whereas cells with the RFTS single mutant (DNMT1^W465A^) and double mutant (DNMT1^W464A/W465A^) exhibited modest and severe impairment of IR resistance (Fig. 4E and F), respectively, confirming the role of DNMT1-mediated DNA methylation in genomic stability maintenance.

Together, the above results strongly indicated that the recognition of H3K9 trimethylation by DNMT1 is important for maintenance DNA methylation, and also genome stability and radiation resistance of ES cells.

## Discussion

The crosstalk between H3K9 methylation and DNA methylation, two of the major epigenetic silencing mechanisms, critically influences gene silencing and heterochromatin formation (17, 18). For instance, previous studies have demonstrated that Suv39h-mediated H3K9-trimethylation promotes the enrichment of DNA methylation at major satellite repeats of pericentromeric heterochromatin (40), which is essential for maintaining heterochromatic assembly and genome stability (41). Whereas the mechanism by which H3K9 methylation and DNA methylation crosstalk has been established in fungi and plants (19, 20), how H3K9me3 is translated into DNA methylation in mammals remains elusive. Through a set of structural, biochemical and cellular analyses, this study shows that the RFTS domain of DNMT1 specifically recognizes H3K9me3 over H3K9me0, in conjunction with the previously identified H3Ub mark. The readout of H3K9me3 not only enhances the stimulation effect of H3Ub on DNMT1 activity but also regulates the genome targeting of DNMT1. This study therefore establishes a direct link between histone H3K9me3 modification and DNMT1-mediated maintenance DNA methylation, which influences the global DNA methylation patterns and genome stability.

Previous studies from others and us demonstrated that DNMT1 assumes autoinhibitory conformations either in the DNA-free state (10, 11) or in the presence of unmethylated CpG DNA (8), in which the autoinhibitory linker located between the CXXC and BAH1 domains serves as a key inhibition-enforcing element in both regulations (8, 10). This study reveals that the recognition between the RFTS domain and H3K9me3 strengthens the RFTS interaction with histone H3 tails that carry either one-or two-mono-ubiquitin mark, directly contributing to the relief of the autoinhibition of DNMT1. The specific recognition of H3K9me3 by the RFTS domains of DNMT1 presumably helps transduce the H3K9me3 signal into DNA methylation, thereby ensuring the epigenetic fidelity of DNA methylation in heterochromatin domains.

DNMT1-mediated maintenance DNA methylation is subjected to a cell cycle-dependent regulation by multiple chromatin regulators, such as UHRF1 (22, 23, 42, 43), Ubiquitin specific protease 7 (USP7) (36, 42–45), PCNA (46, 47) and PAF15 (29, 30). The RFTS-H3K9me3 interaction reinforces the previously identified UHRF1-H3K9me3 axis on chromatin targeting of DNMT1. UHRF1 harbors a tandem TUDOR domain that recognizes H3K9me3 (48–53) and a RING finger domain that mediates H3 ubiquitylation for DNMT1 targeting (25, 26, 28), thereby serving as a platform for the functional crosstalk between H3K9me3 and DNA methylation (54). However, disruption of the interaction between H3K9me3 and UHRF1 via the TUDOR domain mutation only leads to a modest (~10%) reduction of DNA methylation. In this regard, the direct readout of H3K9me3Ub by the RFTS domain of DNMT1 provides a potentially redundant mechanism in transducing H3K9me3 into maintenance DNA methylation. Note that the RFTS-H3K9me3Ub readout does not involves the discrimination of the methylation state of DNA substrates, therefore providing a mechanism in supporting the region-specific methylation maintenance by DNMT1, as opposed to site-specific methylation maintenance (55, 56). Consistently, impairment of this interaction in cells compromises the DNMT1-mediated CpG methylation, leading to an aberrant landscape of DNA methylation and defects in maintenance of genome stability. It remains to be determined whether the DNMT1 mutations introduced in this study also affect the interaction of DNMT1 with other regulatory factors, such as PAF15. This targeting-coupled allosteric stimulation mechanism is reminiscent of the role of histone H3K4me0 in DNMT3A-mediated *de novo* DNA methylation, in which the specific recognition of the DNMT3A ADD domain with H3K4me0 allosterically stimulates its enzymatic activity, thereby providing a mechanism of locus-specific DNA methylation establishment (57).

The DNMT1 RFTS domain adds to the reader modules that offer interpretation of specific histone modifications (34), but deviates from the typical Kme3 readout that depends on aromatic or other hydrophobic residues (34): the RFTS domain presents a single tryptophan to stack against H3K9me3, unlike the typical Kme3 readout involving a hydrophobic cage composed of multiple aromatic residues (34). These observations highlight the evolutionary divergence of the histone modification-binding mechanisms.

## Materials and Methods

### Plasmids

The plasmid that contains DNMT1 was purchased from Addgene (cat # 24952). The DNMT1 cDNA was fused to an N-terminal 3xFlag tag by PCR, followed by subcloning into the pPyCAGIP vector (58) (kind gift of I. Chambers). DNMT1 point mutation was generated by a QuikChange II XL Site-Directed Mutagenesis Kit (Agilent), with the residue numerations based on the isoform 1 of DNMT1 that contains 1616 amino acids. For domain analysis of the DNMT1 RFTS-H3K9me3 interactions, DNA encoding the human DNMT1 RFTS domain (residues 351-597, hDNMT1_RFTS_) or the bovine DNMT1 RFTS domain (residues 349-594, bDNMT1_RFTS_) was cloned into a modified pRSF-Duet vector preceded by an N-terminal His_6_-SUMO tag and ULP1 (ubiquitin-like protease 1) cleavage site. For analysis of hDNMT1_351-1600_ methylation activity, the hDNMT1 construct was inserted into an in-house expression vector as a His_6_-MBP-tagged form. All plasmid sequences were verified by sequencing before use.

### Protein purification

The plasmids were transformed into BL21(DE3) RIL cells (Novagen Inc). When the cell density reached an optical density at 600 nm (OD_600_) of 0.6, the protein expression was induced by 0.1 mM isopropyl β-D-1-thiogalactopyranoside (IPTG) at 16 °C overnight. The cells were harvested, and subsequently resuspended and lysed in a buffer containing 50 mM Tris-HCl (pH 7.5), 25 mM imidazole, 1 M NaCl, 0.5 mM DTT and 1 mM PMSF. The His_6_-SUMO-tagged hDNMT1_RFTS_ and bDNMT1_RFTS_ proteins were first purified using a nickel column with elution buffer containing 25 mM Tris-HCl (pH 8.0), 100 mM NaCl and 300 mM imidazole. The eluted protein was incubated with ULP1 on ice for cleavage of the His_6_-SUMO tag, followed by purification of the tag-free protein by anion exchange chromatography on a HiTrap Q XL column (GE Healthcare) and nickel affinity chromatography. The protein sample was finally purified on size-exclusion chromatography on a HiLoad 16/600 Superdex 75 pg column (GE Healthcare), pre-equilibrated with buffer (20 mM Tris, pH 7.5, 50 mM NaCl, 5 mM DTT). The His_6_-MBP tagged DNMT1 proteins were first purified by Ni^2+^ chromatography, followed by ion-exchange chromatography on a Heparin HP (GE Healthcare) or Q HP column (GE Healthcare), removal of His_6_-MBP tag by TEV protease cleavage, a second round of nickel affinity chromatography, and size-exclusion chromatography on a Superdex 200 16/600 column (GE Healthcare). DNMT1 mutants were introduced by site-directed mutagenesis and purified as that described for wild-type protein. All purified protein samples were stored at −80 °C before use.

### Chemical modifications of histones

To prepare the unbiquitylated histones, the His_6_-SUMO-Ub(G76C) and each synthesized histone peptide (H3_1-25_K18C, H3_1-25_K9me3K18C, H3_1-24_K18CK23C or H3_1-_ 24K9me3K18CK23C, each containing an additional tryptophan at the C-terminus), dissolved in buffer (250 mM Tris-HCl (pH 8.6), 8 M urea, 5 mM TCEP), were mixed in a 4:1 molar ratio, and incubated at room temperature for 30 min. The crosslinker 1,3-dichloroacetone, dissolved in N,N’-dimethylformamide, was added to the reaction mixture with the amount equal to one-half of the total sulfhydryl groups. After 2 hr.-incubation on ice, the reaction was stopped by 5 mM β-ME. Purification of H3_1-25_K18Ub, H3_1-25_K9me3K18Ub, H3_1-24_Ub2 and H3_1-24_K9me3Ub2 were achieved through cation-exchange chromatography on a mono S column (GE Healthcare).

### Crystallization and structure determination

To crystallize the bDNMT1_RFTS_-H3K9me3-Ub2 complex, bDNMT1_RFTS_ was mixed with H3_1-24_K9me3K18CK23C peptide, containing an additional tryptophan at the C-terminus, and ubiquitin in a 1:1:2 molar ratio. The complex was incubated on ice for 30 min before crystallization. The crystals were generated in a buffer containing 0.1 M citric acid (pH 3.5), 28% PEG8000 at 4 °C, using the hanging-drop vapor diffusion method. The crystals were soaked in the crystallization buffer supplemented with 20-25% (v/v) glycerol as cryo-protectant before flash frozen in liquid nitrogen. The X-ray diffraction data were collected on the beamline BL5.0.1 at Advanced Light Source (ALS), Lawrence Berkeley National Laboratory. The data were indexed, integrated and scaled by HKL2000 program (59) or XDS(60). The structure was solved by molecular replacement using PHASER(61) with the RFTS domain in the DNMT1 structure (PDB 4WXX) as searching model. Iterative cycles of model rebuilding and refinement were carried out using COOT (62) and PHENIX (63), respectively. Data collection and structure refinement statistics were summarized in Table S2.

### ITC binding assay

ITC measurements were performed using a MicroCal iTC200 instrument (GE Healthcare). Synthesized H3_1-22_, H3_1-22_K9me3, H3_1-22_K9Ac, H3_1-15_K4me3, H3_21-_ 33K27me3, H331-43K36me3 and H414-25K20me3 peptides each contain a C-terminal tyrosine for spectroscopic measurement. To measure the bindings between hDNMT1_RFTS_ and the peptides, 1 mM peptide was titrated with 0.1 mM hDNMT1_RFTS_ at 20 °C. To measure the bindings between hDNMT1_351-1600_ and H3Ub2 or H3K9me3Ub2 peptides, 0.12 mM peptide was titrated with 12 μM hDNMT1_351-1600_ sample at 5 °C. Prior to the titration, both peptide and protein samples were subjected to overnight dialysis against buffer containing 20 mM Tris-HCl (pH 7.5),100 mM NaCl and 1 mM DTT. Buffer to buffer titration was performed to ensure no abnormality of base line. Analyses of all data were performed with MicroCal Origin software, fitted with single-site binding mode. The ITC parameters were summarized in Table S1.

### DNA methylation kinetics assay

The DNA methylation assays were performed as previously described (8) with modifications. Synthesized H3_1-25_, H3_1-25_K9me3, H3_1-25_K18Ub, H3_1-25_K9me3K18Ub, H31-24Ub2 and H31-24K9me3Ub2 peptides, each with a C-terminal tryptophan, were used for evaluation of the RFTS-mediated enzymatic stimulation of hDNMT1351-1600. Each reaction mixture contains 0.1 μM hDNMT1_351-1600_, wild type or mutants, 0.5 mM S-adenosyl-L-[methyl^3^H] methionine (SAM) (Perkin Elmer), 0.4 μM (GT^m^C)_12_/(GAC)_12_ hemimethylated DNA duplex, and various amount of histone peptides in 50 mM Tris-HCl (pH 8.0), 7 mM β-ME, 5% glycerol, 100 μg/mL BSA and 100 mM NaCl, unless indicated otherwise. The reaction mixture was incubated at 37 °C for 20 min, before quenched by 2 mM cold SAM. Eight μL of the reaction mixture was applied onto DEAE filtermat (Perkin Elmer), sequentially washed with 0.2 M ammonium bicarbonate (twice), water and ethanol. The filter paper was then air dried and soaked in ScintiVerse cocktail (Thermo fisher). The activity was measured by Beckman LS6500 scintillation counter.

### Electrophoretic mobility shift assay

To measure the H3_1-24_K9me3Ub2-binding affinity, 1 μM hDNMT1_351-1600_ or hDNMT1_RFTS_ protein was incubated with various amount of H31-24K9me3Ub2 peptide in 10 μL binding buffer (20 mM Tris–HCl (pH 7.5), 100 mM NaCl, 1 mM DTT and 5% glycerol) at 4 °C for 1 hr. The sample mixture was resolved in 4-10% native gel using 0.5X TG buffer at 4 °C under 100 V for 3.5 hr. The gel image was visualized by coomassie blue staining.

To measure the DNA-binding affinity of hDNM1_351-1600_, 0.1 μM 26-base pair DNA duplex containing one central hemimethylated CpG site (upper strand: 5’-ACACCAAGCCTGMGGAGGCTCACGGA-3’, M = 5-methylcytosine; lower strand: 5’-TCCGTGAGCCTCCGCAGGCTTGGTGT-3’) was mixed with 0, 1, 2 or 5 ìM hDNMT1351-1600, wild type or mutants, in the presence or absence of the H31-24K9me3Ub2 peptide, in buffer containing 20 mM Tris–HCl (pH 7.5), 50 mM NaCl, 1 mM DTT and 5% glycerol at 4 °C for 1 hr, before resolved in a 6% TBE native gel. The protein-DNA complex was visualized by SYBR green staining.

### Cell lines and tissue culture

Dnmt1-knockout mouse embryonic stem cells (1KO-ESCs; a gift from Dr. M. Okano) were cultivated as previously described on gelatin-coated dishes in the base ESC culture medium supplemented with leukemia inhibitory factor(64). 1KO-ESCs were transfected by Lipofectamine 2000 (Invitrogen) with the pPyCAGIP empty vector or that carrying WT or mutant DNMT1. Forty-eight hours post-transfection, the transduced ES cells were selected out in culture medium with 1μg/mL puromycin for over two weeks. The pooled stable-expression cell lines and independent single-cell-derived clonal lines were first established, followed by further characterizations such as immunoblotting of DNMT1.

### Antibodies and Western blotting

Antibodies used for immunoblotting include α-Flag (Sigma; M2), H4K20me3 (Abcam ab9053), H3K9me3 (Abcam ab8898), general histone H3 (Abcam ab1791) and α-Tubulin (Sigma). Whole cell protein lysates were prepared by NP40 lysis buffer (50mM Tris-HCl pH 8.0, 150mM NaCl, 1% NP-40) followed by brief sonication and centrifugation. After mixing with loading buffer and boiling for 5 minutes, the same amount of extracted samples were loaded into SDS-PAGE gels for immunoblotting analysis as previously described (64, 65).

### Quantification of 5-methyl-2’-deoxycytidine (5mC) in genomic DNA

Enzymatic digestion of cellular DNA and LC-MS/MS measurements of the levels of 5-mdC in the resulting nucleoside mixture was carried out as described previously (64).

### Confocal immunofluorescence (IF)

The immunofluoresence was carried out as described before (66). In brief, 3xFlag-tagged DNMT1 transduced cells were fixed with 4% paraformaldehyde and 10-min permeabilization in 0.1% Triton X-100. After one-hour block in PBS with 2.5% of BSA, cells were stained with primary antibodies, followed by staining with the Alexa-488 or Alexa-594 conjugated secondary antibodies. The primary antibodies used include M2 anti-FLAG (Sigma) and H3K9me3 (Abcam). Fluorescence was detected in a FV1000 confocal microscope (UNC Imaging Core).

### Co-immunoprecipitation (CoIP)

Cells were lysed in NP40 lysis buffer that contains 50mM Tris-HCl (pH 8.0), 150mM NaCl, 1% of NP-40, 1mM DTT and a complete protease inhibitor (Roche) as described (66, 67). Antibodies H4K20me3 (Abcam ab9053), H3K9me3 (Abcam ab8898) and general histone H3 (Abcam ab1791) conjugated with protein A/G beads (Millipore) or anti-Flag M2-conjugated agarose beads (Sigma) were incubated with the lysates overnight at 4 °C. The beads were then washed 3–6 times with cell lysis buffer, and the bound proteins were eluted in SDS buffer and analyzed by western blotting.

### Preparation of enhanced reduced representation bisulfite sequencing (eRRBS) libraries

Construction of eRRBS libraries was performed as described before (64). Briefly, 1 microgram of total genomic DNA (gDNA) was added with 0.1% of unmethylated lambda DNA (Promega), followed by one-hour digestion with three enzymes (MspI, MseI and BfaI) at 37 degree. The purified digested gDNA was then subjected to end repair, A-tailing and ligation to NEBNext Methylated Adaptors (NEBNext DNA Library Prep Kit), followed by purification using AMPure beads. Bisulfite conversion and library construction were carried out as before using the EpiMark Bisulfite Conversion Kit (NEB cat# E3318) according to manufacturer’s specifications. The generated multiplexed eRRBS libraries were subjected to deep sequencing in an Illumina HiSeq 4000 platform with a paired end PE150 cycle (carried out by UNC HTSF Genomic Core).

### eRRBS data processing

General quality control checks were performed with FastQC v0.11.2 (http://www.bioinformatics.babraham.ac.uk/projects/fastqc/). The last 5 bases were clipped from the 3’ end of every read due to questionable base quality in this region, followed by filtration of the sequences to retain only those with average base quality scores of more than 20. Examination of the 5’ ends of the sequenced reads indicated that 70%-90% (average 83.0%) were consistent with exact matches to the expected restriction enzyme sites (i.e. MspI, BfaI, and MseI). Approximately 85% of both ends were consistent with the expected enzyme sites. Adapter sequence was trimmed from the 3’ end of reads via Cutadapt v1.2.1 (parameters -a AGATCGGAAGAG -O 5 -q 0 -f fastq; https://doi.org/10.14806/ej.17.1.200). Reads less than 30nt after adapter-trimming were discarded. Filtered and trimmed datasets were aligned via Bismark v0.18.1 (parameters -X 1000 --non_bs_mm)(68), using Bowtie v1.2(69) as the underlying alignment tool. The reference genome index contained the genome sequence of enterobacteria phage λ (NC_001416.1) in addition to the mm10 reference assembly (GRCm38). For all mapped read pairs, the first 4 bases at the 5’ end of read1 and the first 2 bases at the 5’ end of read2 were clipped due to positional methylation bias, as determined from QC plots generated with the ‘bismark_methylation_extractor’ tool (Bismark v0.18.1). To avoid bias in quantification of methylation status, any redundant mapped bases due to overlapping read ends from the same read pair were trimmed. Read pairs in which either read end had 3 more or methylated cytosines in non-CpG context were assumed to have escaped bisulfite conversion and were discarded. Finally, mapped read pairs were separated by genome (mm10 or phage λ). Read pairs mapped to phage λ were used as a QC assessment to confirm that the observed bisulfite conversion rate was >99%. Read pairs mapped to the mm10 reference genome were used for downstream analysis.

### eRRBS data analysis

Although the eRRBS data does carry stranded information, data from the plus and minus strands have been collapsed in this analysis. Weighted methylation scores were calculated as described by Schultz et al(70) at query regions of interest, such as H3K9me3 and H4K20me3 peaks. Depth tracks for the eRRBS data at all genomic loci were first generated in bedGraph format by BEDTools v2.24.0(71) ‘genomecov’, then converted to bigWig format by bedGraphToBigWig (UCSC utility script; http://hgdownload.soe.ucsc.edu/admin/exe/). Percent methylation tracks for the eRRBS data at CpG sites in non-cytosine context were generated by first constructing WIG tracks reporting %methylation (count of methylated C bases / total C bases) per site, followed by conversion to bigWig format by wigToBigWig (UCSC utility script).

### Neutral comet assay

Neutral comet assays were performed using the CometAssay Reagent Kit (Trevigen) according to the manufacturer’s instruction. Cells were plated and treated with ionizing radiation (IR; 5Gy). After 2hr post-IR treatment, cells were harvested and mixed with LMAgarose (Trevigen). The mixtures were placed on glass slides (Trevigen). Cells were lysed with lysis solution (Trevigen) at 4 °C for 1h. The slides were washed with TBE (90mM Tris borate (pH 8.3) to removal of remain lysis solution and subject to electrophoresis at 40V for 40 min in TBE buffer. Samples were fixed with 70% ethanol for 10 min. The DNA was stained with SYBR-green (Invitrogen) for 15 min in RT. The images were taken by fluorescence microscopy and tail moments were calculated for at least 100 cells for each sample by Image J (v 1.48). Tail moment (TM) reflects both the tail length (TL) and the fraction of DNA in the comet tail (TM = %DNA in tail × TL/100).

### Statistics

The two-tailed Student t-tests were performed to compare distributions between different groups. And the p value lower than 0.01 was considered to be statistically significant.

### Accession codes

Coordinates and structure factors for the bDNMT1RFTS-H3K9me3-Ub2 complex have been deposited in the Protein Data Bank under accession codes 6PZV. The eRRBS data have been deposited in Gene Expression Omnibus (GEO) under accession code GSE145698.

## Acknowledgments

We would like to thank staff members at the Advanced Light Source (ALS), Lawrence Berkeley National Laboratory for access to X-ray beamlines. We are also grateful for professional support of UNC facilities including Genomics Core, which are partly supported by the UNC Cancer Center Core Support Grant P30-CA016086. This work was supported by NIH grants (1R35GM119721 to J.S, 5R21ES025392 to Y.W., RO1GM079641 to O.G., 1R35GM126900 to B.D.S., 1R01CA215284 and 1R01CA211336 to G.G.W., and CA198279 and CA201268 to K.M.M.), NIH T32 Training Fellowship for Integrated Training in Cancer Model Systems (T32CA009156) and an American Cancer Society Postdoctoral Fellowship (PF-19-027-01–DMC) to C.J.P. and grants from When Everyone Survives (WES) Leukemia Research Foundation (to G.G. W.) and Gabrielle’s Angel Foundation for Cancer Research (to G.G.W.). D.J.P. was supported by a SCOR grant by the Leukemia and Lymphoma Society and by the Memorial Sloan-Kettering Cancer Center Core Grant (P30CA008748). This work was also supported in part by the Intramural Research Program of the National Institute of Environmental Health Sciences, NIH (ES101965 to PAW). J.P.C. is supported in part by T32 CA09302. G.G.W. and K.M.M. are American Cancer Society (ACS) Research Scholar, C.J.P. is an ACS postdoctoral fellow, and G.G.W. is a Leukemia & Lymphoma Society (LLS) Scholar.

## Author Contributions

W.R., H.F., S.A.G., Y.G., J.J.K., L.L., C.J.P., X-F. T., Z-M. Z., J.P.C, J.Y., L.G., S.L., L.C., B.D., Y.W., Q.C., and S.O. performed experiments. Y.W., B.S., O.G., K.M.M., S.O., P.A.W., D.J.P, G.G.W. and J.S. organized the study. D.J.P., O.G., G.G.W. and J.S. conceived the study, G.G.W. and J.S wrote the manuscript with input from all the authors.

## Supplementary Information for

**Figure S1.**
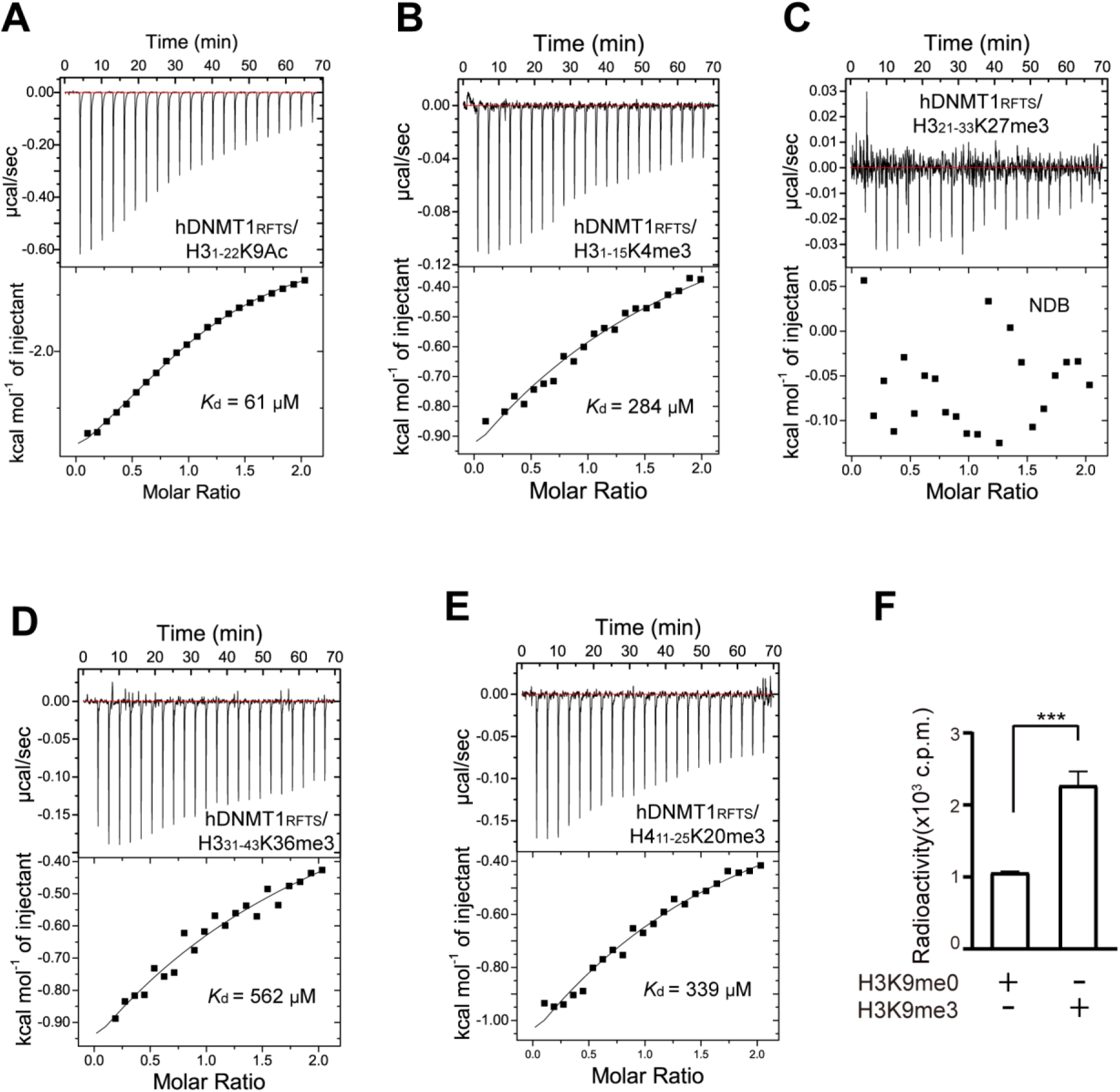
Biochemical analysis of the interaction between DNMT1 RFTS domain and histone peptides. (**A-E**) ITC binding curves of hDNMT1_RFTS_ with the H3_1-22_K9Ac peptide (**A**), H3_1-15_K4me3 peptide (**B**), H3_21-33_K27me3 peptide (**C**), H3_31-41_K36me3 peptide (**D**) and H4_11-25_K20me3 peptide (**E**). (**F**) DNA methylation activity of hDNMT1_351-1_6_00_ in the presence of 100-fold molar excess of H3K9me0 or H3K9me3 peptides. Mean and s.d. were derived from three independent measurements. (***, *p* < 0.001, Student’s *t*-test).

**Figure S2.**
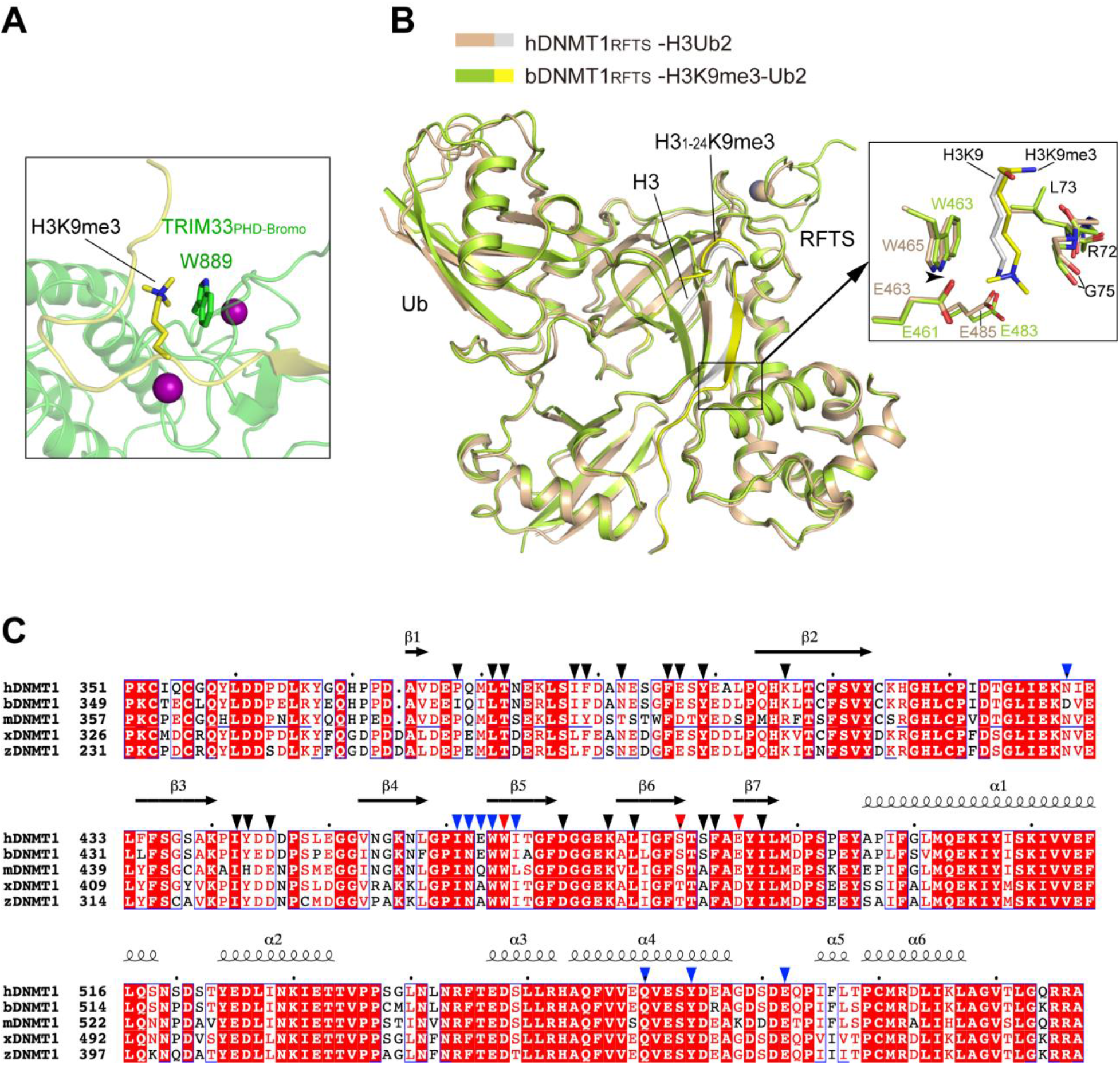
Structural analysis of the H3K9me3-binding pocket. (**A**) Close-up view of the interaction between the residue W889 of the TRIM33 PHD domain and H3K9me3 (PDB 3U5N). (**B**) Structural superposition of bDNMT1_RFTS_-H3K9me3-Ub2 and hDNMT1_RFTS_-H3Ub2 complexes (PDB 5WVO), with the H3K9me3-binding sites shown in the expanded view. (**C**) Sequence alignment of the DNMT1 RFTS domain from human (hDNMT1), bovine (bDNMT1), mouse (mDNMT1), Xenopus Laevis (xDNMT1) and Zebrafish (zDNMT1). Strictly conserved residues are colored in white and shaded in red. Similar residues are colored in red. The secondary structures corresponding to hDNMT1 RFTS are marked on top. The residues for interaction with H3K9me3 peptide are marked by blue arrows. The residues for interacting with ubiquitin molecules are marked by dark arrows. The H3 K9me3-interacting residues are marked by red arrows.

**Figure S3.**
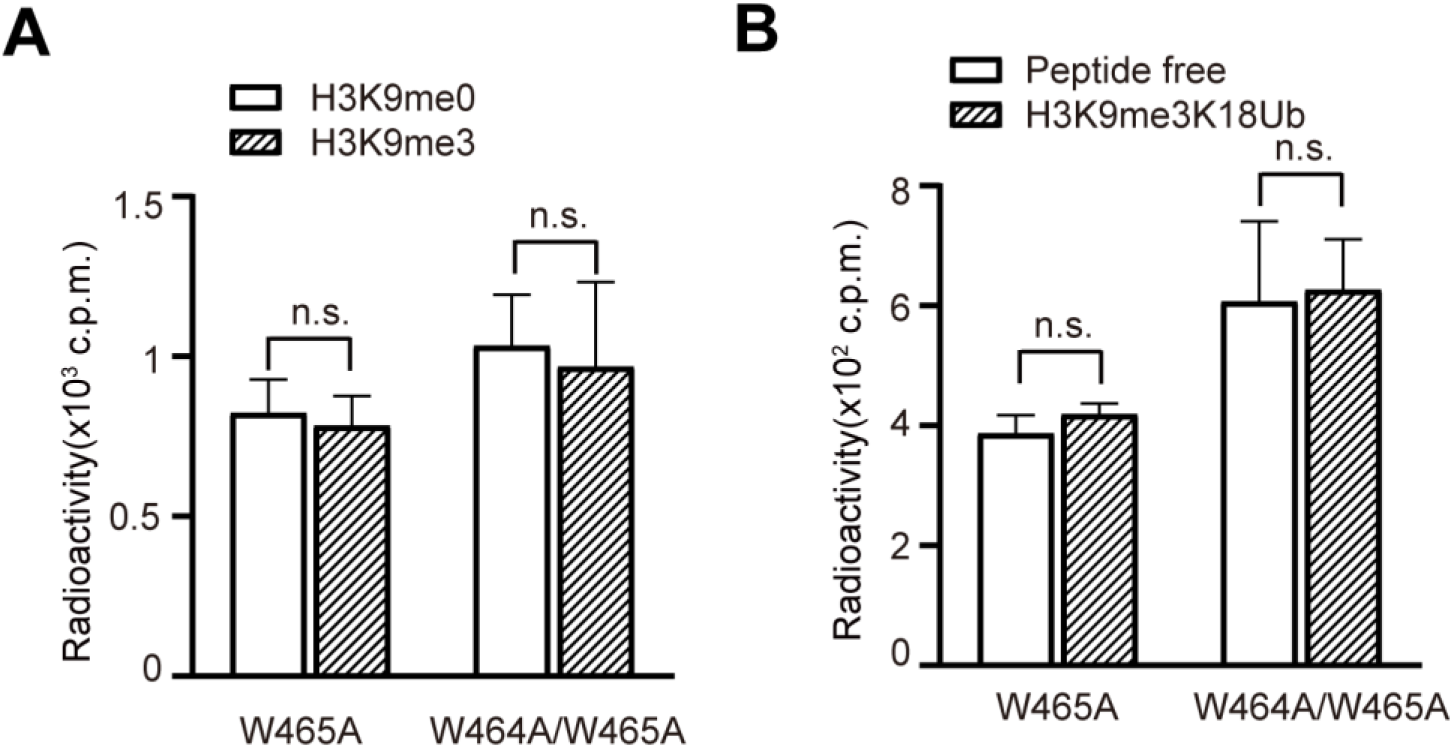
*In vitro* DNA methylation assays of hDNMT1_351-1600_ mutants with H3 peptide. (**A**) DNA methylation activity of W465A-or W464A/W465A-mutated hDNMT1_351-1600_ in the presence of H3K9me0 or H3K9me3 peptides. (**B**) DNA methylation activity of W465A- or W464A/W465A-mutated hDNMT1_351-1600_ in the absence or presence of the H3K9me3Ub. Mean and s.d. were derived from three independent measurements. (n.s. not significant).

**Figure S4.**
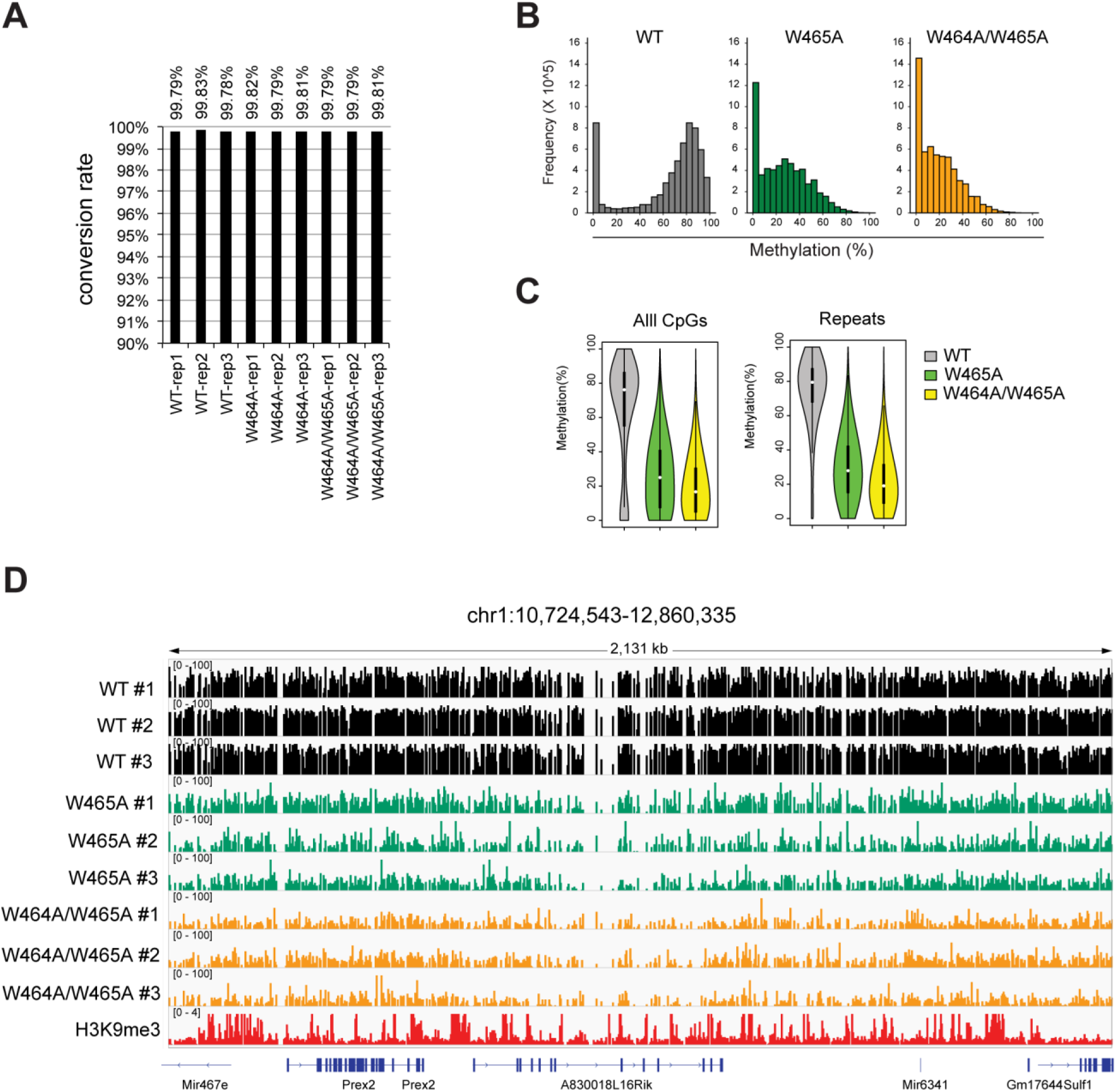
eRRBS profilings of DNA methylation in 1KO-ESC cells reconstituted with either WT or RFTS-mutated DNMT1. (**A**) The rates of bisulfite conversion, labeled on top of columns, for all cytosines in each 1KO-ESC cell sample with expression of the indicated DNMT1, as determined by the unmethylated lambda DNA used as spike-in controls. (**B**) Distribution of absolute methylation levels for CpG sites with >5 coverage among 1KO-ESC lines with stable expression of the indicated DNMT1, as detected by eRRBS. (**C**) Violin plots showing distribution of absolute methylation levels for CpG sites with >5 coverage at all CpG sites or those within the repeated genomic sequences among the indicated cells samples. White dots are the median and box lines are the first and third quartile of the data. (**D**) Representative IGV views of CpG methylations at an H3K9me3 (red, bottom)marked genomic region located in the chromosome 1 among three replicated 1KO-ESC lines with stable expression of the indicated DNMT1. Cytosines covered by at least 5 reads according to eRRBS data are shown, with each site designated by a vertical line.

**Table S1.** Summary of ITC binding parameters.

| Protein | Peptide | $K_d$ (μM) | N value |
| --- | --- | --- | --- |
| hDNMT1<sub>RFTS</sub>, WT | H3(1-22)<sup>#</sup> | 6.4 ± 0.8 | 0.74±0.05 |
| hDNMT1<sub>RFTS</sub>, WT | H3(1-22)K9me3<sup>#</sup> | 1.3 ± 0.1 | 0.77± 0.1 |
| hDNMT1<sub>351-1600</sub>, WT | H3(1-24)Ub2<sup>#</sup> | 0.091 ± 0.02 | 0.91± 0 |
| hDNMT1<sub>351-1600</sub>, WT | H3(1-24)K9me3Ub2<sup>#</sup> | 0.017 ± 0.003 | 0.96±0.1 |
| hDNMT1<sub>RFTS</sub>, WT | H3(1-22)K9Ac | 61 ± 14 | 0.95±0.1 |
| hDNMT1<sub>RFTS</sub>, WT | H3(1-15)K4me3 | 284 ± 32 | 1.0* |
| hDNMT1<sub>RFTS</sub>, WT | H3(21-33)K27me3 | NDB |  |
| hDNMT1<sub>RFTS</sub>, WT | H3(31-43)K36me3 | 562 ± 77 | 1.0* |
| hDNMT1<sub>RFTS</sub>, WT | H4(14-25)K20me3 | 339 ± 11 | 1.0* |
| hDNMT1<sub>RFTS</sub>, W465A | H3(1-22)K9me3 | 22 ± 0.7 | 1.1±0.04 |
| hDNMT1<sub>RFTS</sub>, W465A | H3(1-22) | 30 ± 6 | 1.0±0.2 |
| hDNMT1<sub>RFTS</sub>, W464AW465A | H3(1-22)K9me3 | NDB |  |

NDB, no detectable binding. \*The N value was set manually. <sup>#</sup>The mean value and S.D. were derived from two-independent measurements.

**Table S2.** Data collection and refinement statistics.

|  |  |
| --- | --- |
|  | bDNMT1 RFTS –<br>Ubiquitin – H3K9me3<br>(PDB: 6PZV) |
| **Data collection** |  |
| Space group | P 2<sub>1</sub> 2<sub>1</sub> 2 |
| Cell dimensions |  |
| *a*, *b*, *c* (Å) | 70.3, 196.4, 67.7 |
| $\alpha$, $\beta$, $\gamma$ (°) | 90, 90, 90 |
| Resolution (Å) | 49.11-3.01(3.12-3.01)<sup>a</sup> |
| *R*<sub>merge</sub> | 0.126(1.023) |
| *I*/$\sigma$(*I*) | 12.3(1.3) |
| *CC*<sub>1/2</sub> | 0.998(0.583) |
| Completeness (%) | 99.1(92.3) |
| Redundancy | 9.8(7.9) |
| **Refinement** |  |
| No. reflections | 19,094 |
| *R*<sub>work</sub> / *R*<sub>free</sub> | 0.212/0.275 |
| No. atoms |  |
| Protein | 6184 |
| Zn<sup>2+</sup> | 2 |
| Water | 48 |
| *B* factors (Å<sup>2</sup>) |  |
| Protein | 85.1 |
| Zn<sup>2+</sup> | 82.0 |
| Water | 67.5 |
| r.m.s deviations |  |
| Bond lengths (Å) | 0.003 |
| Bond angles (°) | 0.588 |

<sup>a</sup>Values in parentheses are for highest-resolution shell.

**Legend for Table S3: Summary of eRRBS data analysis.**

## Notes

### Competing Interest Statement

O.G. and B.D.S are co-founders of EpiCypher Inc.

